# MutLγ enforces meiotic crossovers in *Arabidopsis thaliana*

**DOI:** 10.1101/2024.09.18.613675

**Authors:** Stéphanie Durand, Qichao Lian, Victor Solier, Joiselle Blanche Fernandes, Raphael Mercier

## Abstract

During meiosis, each chromosome pair experiences at least one crossover (CO), which directs their balanced segregation in addition to shuffling genetic information. COs tend to be away from each other, a phenomenon known as CO interference. The main biochemical pathway for CO formation, which is conserved in distant eukaryotes, involves the ZMM proteins together with the MLH1-MLH3 complex (MutLγ). Here, we aim to clarify the role of MutLγ in CO formation in *Arabidopsis thaliana*. We show that AtMutLγ is partially dispensable for ZMM-dependant CO formation. HEI10 large foci - that mark CO sites in wild-type-form at a normal level in *mlh1* and *mlh3* mutants, but are inefficiently maturated into COs. Mutating the *MUS81* nuclease in either *mlh1* or *mlh3* leads to chromosome fragmentation, which is suppressed by further mutating the *zmm msh5*. This suggests that in the absence of MutLγ, recombination intermediates produced by ZMMs are resolved by MUS81, which does not ensure CO formation. Finally, CO interference is not affected in *mlh1*, which is compatible with a random sub-sampling of normally patterned CO sites. We conclude that AtMutLγ imposes designated recombination intermediates to be resolved exclusively as COs, supporting the view that MutLγ asymmetrically resolves double-Holliday junctions, yielding COs.

## Introduction

Meiotic recombination initiates with the formation of a large number of DNA double-strand breaks followed by strand exchange with the homologous chromosomes forming DNA joint-molecule intermediates. A subset of these joint molecules is matured into double Holliday junctions (dHJs), two adjacent branched DNA structures that contain four double-stranded arms (1–3). The formation of dHJs is promoted by a group of evolutionarily conserved proteins collectively named ZMMs (originally for the yeast Zip1-4, Msh4-5, Mer3) (4). Holliday junctions are symmetrical and their resolution can in principle lead to both crossovers (COs) and non-crossovers (NCOs), but, at least in budding yeast, they are processed almost exclusively as COs by the MLH1/MLH3 (MutLγ) complex (5,6). In *vitro* studies with mammalian proteins confirmed the capacity of MutLγ to cleave DNA, an activity promoted by EXO1 and PCNA, and suggested a mechanism for the biased processing of dHJs into COs (7,8). The COs promoted by the ZMMs and MutLγ, are called class I COs. A second minor pathway is independent of ZMMs and involves structure-specific DNA endonuclease including MUS81 (9).

In Arabidopsis, the ZMM proteins are responsible for most COs, with a ∼90% drop in CO formation in the absence of any of them (10,11). The MLH1 and MLH3 proteins accumulate in bright foci at future class I CO sites at late pachytene and persist until diakinesis (12–14). Accordingly, MLH1 foci are absent at diakinesis in the *zmm* mutants *hei10* (*Zip3* homolog) and *zip4* (12,15). Disruption of *MLH3* leads to the loss of about half of the meiotic crossovers, a milder defect than a *zmm* mutant, associated with the loss of the obligate crossover and the presence of univalents (13). Mutants in *MLH1* have fertility defects, but to our knowledge, the meiotic phenotype was not described (16). Here we aimed to clarify the function of MutLγ in Arabidopsis meiosis and we conclude that the function of AtMutLγ is to impose ZMM recombination intermediates to be resolved exclusively as COs, which is compatible with the proposed biochemical function in processing dHJs asymmetrically.

## Results and discussion

### MLH1 and MLH3 are partially dispensable for class I CO formation

The *zmm* mutants in Arabidopsis *msh4, msh5, shoc1, ptd1, hei10,* and *zip4* (homologs of the yeast *msh4, msh5, zip2, spo16, zip3 and zip4*, respectively) exhibit a ∼85% reduction in CO formation leading to the presence on average of ∼3 pairs of univalents (among 5 chromosome pairs) (15,17–21) (Figure 1). The residual crossovers in *zmm* mutants are attributed to the class II pathway, which acts in parallel to the *zmm* pathway. Accordingly, combining *zmm* mutations does not reduce further CO formation (22). We performed chromosome spreads on male meiocytes of *mlh1*, *mlh3,* and the *zmm* representative *msh5* (Figure 1). Compared to *msh5*, and other previously characterized *zmm* mutants, both *mlh1* and *mlh3* mutants have a moderate reduction in meiotic CO formation leading to the formation of only ∼1.5 univalents (p<10^-6^, Figure 1E) (13,22). Combining *mlh1* and *mlh3* mutations did not increase the frequency of univalents (p=0.20, Figure 1), suggesting that MLH1 and MLH3 act together to promote CO formation. Consistently, MLH1 and MLH3 form co-foci (13), and MLH1 foci formation depends on *mlh3* (Figure 1F-G, n=22 diplotene/diakinesis cells). The combination of *mlh3* with *msh5* or with *shoc1* results in a *zmm*-like level of univalent (22). Similarly, the *mlh1 msh5* mutant is indistinguishable from the *msh5* single mutant in terms of univalent frequency (p=0.78, Figure 1E) and is more affected than *mlh1* (p<10^-6^). This shows that residual class I COs are formed in the absence of MLH1 or/and MLH3. Altogether, this demonstrates that MLH1 and MLH3 act together in the class I/ZMM pathway but with a less essential role than the ZMMs for CO formation.

**Figure 1:**
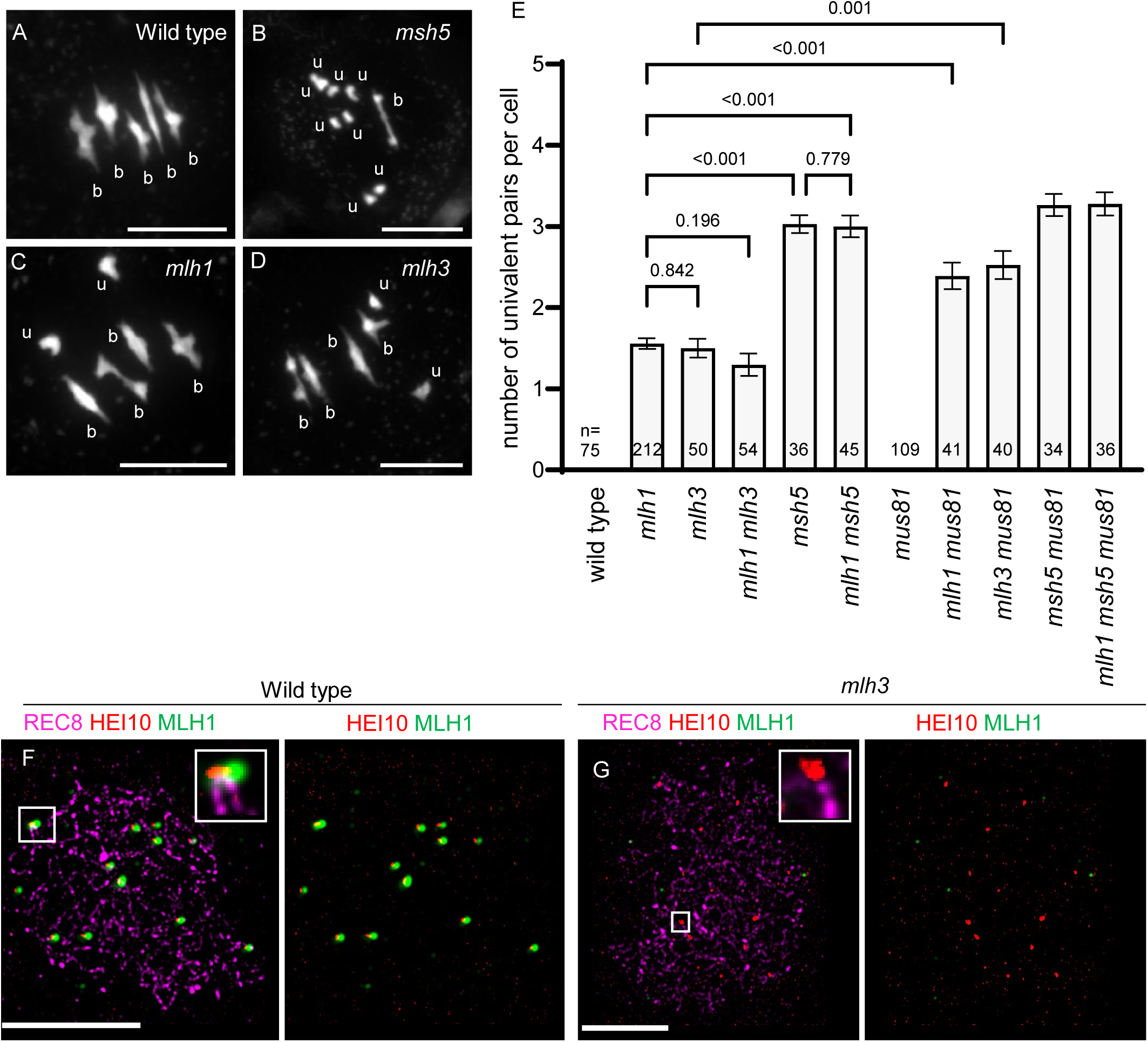
MLH1 and MLH3 are partially dispensable for class I CO formation. (A-D) Metaphase I chromosome spreads for both wild-type and mutant male meiosis, stained with DAPI. Pairs of homologous chromosomes connected by crossovers are referred to as bivalents, abbreviated as “b.” When crossovers are absent and homologous chromosomes are separated, they are referred to as univalents, abbreviated as “u”. Scale bar=10µm. (E) Quantification of the number of univalent pairs per male meiotic cell. If zero pairs of univalents were observed, the bar is not visible. The mean +/-SEM is shown and the number n of analyzed cells is indicated. P values are Kuskal-Wallis and uncorrected Dunn’s test performed in Prism 10.2.0. (F-G) Immunolocalization of REC8 (purple), HEI10 (Red) and MLH1 (green) at diakinesis of male meiocytes. (F) In wild type, MLH1 and HEI10 colocalize in foci. (G) In *mlh3*, HEI10 foci are present (see also Figure 3), but MLH1 foci are not detected. Scale bar=3µm.

## In absence of MLH1 or MLH3, MUS81 becomes crucial for CO resolution

The yeast and mammal MLH1/MLH3 complex is proposed to resolve double holiday-junctions (dHJs) in a biased manner resulting exclusively in COs (7). In the absence of bias, dHJs resolution is predicted to result in 50% of COs and 50% of NCOs. We suggest that in the absence of Arabidopsis MLH1/MLH3, dHJs are resolved in an unbiased manner, leading to the loss of ∼50% of class I COs, explaining the moderate CO reduction in *mlh1* and *mlh3* compared to *zmm* mutants. A candidate for supporting this unbiased activity is the structure-specific nuclease MUS81, which is involved in class II CO formation (23). MUS81 has only a minor role in supporting CO formation, and no univalents are observed in *mus81* mutants (Figure 1E, n=109). However, mutating *MUS81* in the *mlh1* or *mlh3* background resulted in an increased frequency of univalents (Figure 1E). Furthermore, *mlh1 mus81* and *mlh3 mus81* double mutants showed chromosome fragmentations from anaphase I onward (Figure 2). This indicates a failure in the resolution of some recombination intermediates when both MLH1/MLH3 and MUS81 are absent. Importantly, chromosome fragmentation in *mlh1 mus81* and *mlh3 mus81* is suppressed by mutating *msh5* (Figure 2), supporting the conclusion that the recombination intermediates produced by the ZMM pathway (dHJs) fail to be repaired if both MLH1/MLH3 and MUS81 are absent.

**Figure 2:**
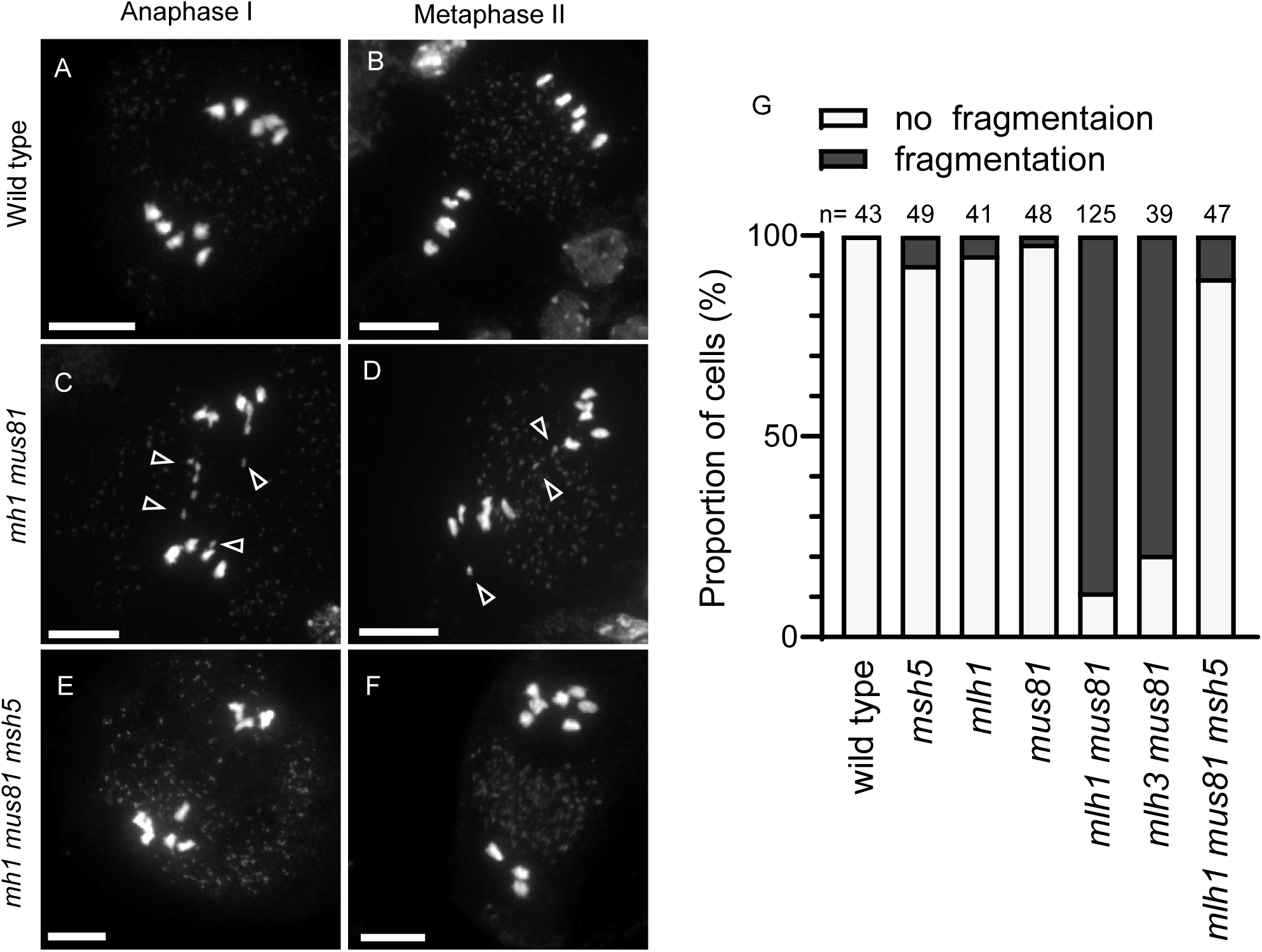
MUS81 becomes crucial for Resolving Crossovers in the Absence of MutLγ. Representative images of Anaphase I and Metaphase II chromosome spreads from wild-type (A,B) and mutant *mlh1 mus81* (C,D) and *mlh1 mus81 msh5* (E,F). Arrowheads indicate chromosome fragments. (G) Percentage of cells in which chromosome fragmentation was detected (black) or not (light grey). The numbers above the bar plots indicate the total number of cells analyzed for each genotype. Scale bar=10µm.

### MLH1 and MLH3 act downstream of HEI10

In the wild type, the HEI10 protein initially decorates the center of the synaptonemal complex, a large zipper-like structure that associates the homologous chromosome all along their length (24) and thus appears as a dotted line in between the two homologous axes while they associate (marked by the REC8 Cohesin on Figure 3A). HEI10 then progressively accumulates in a limited number of large foci that co-localize with MLH1 (Figure 3 B-C). MLH1-HEI10 co-foci mark the sites of COs at the end of pachytene, diplotene and diakinesis in wild type (Figure 1F). It is proposed that the dynamic of HEI10 – its coarsening-may drive the selection of CO sites (14,15,25,26). In *mlh1* and *mlh3*, we observed the same initial HEI10 dynamic as in the wild type (Figure 3 D-I): Initially, numerous HEI10 small foci decorate the center of the synaptonemal complex along its entire length (Figure 3D, 3G). With meiosis progression, HEI10 gradually forms larger and less numerous foci, culminating in ∼12 HEI10 foci at diplotene, indistinguishably in wild-type, *mlh1* and *mlh3* (Figure 3B, 3E, 3H, 3J). However, while the number of HEI10 foci is stable from diplotene to diakinesis in the wild type, their number drops dramatically in *mlh1* and *mlh3* (Figure 3C, 3F, 3I, 3J). This suggests that MutLγ stabilizes the HEI10 foci in late prophase.

**Figure 3:**
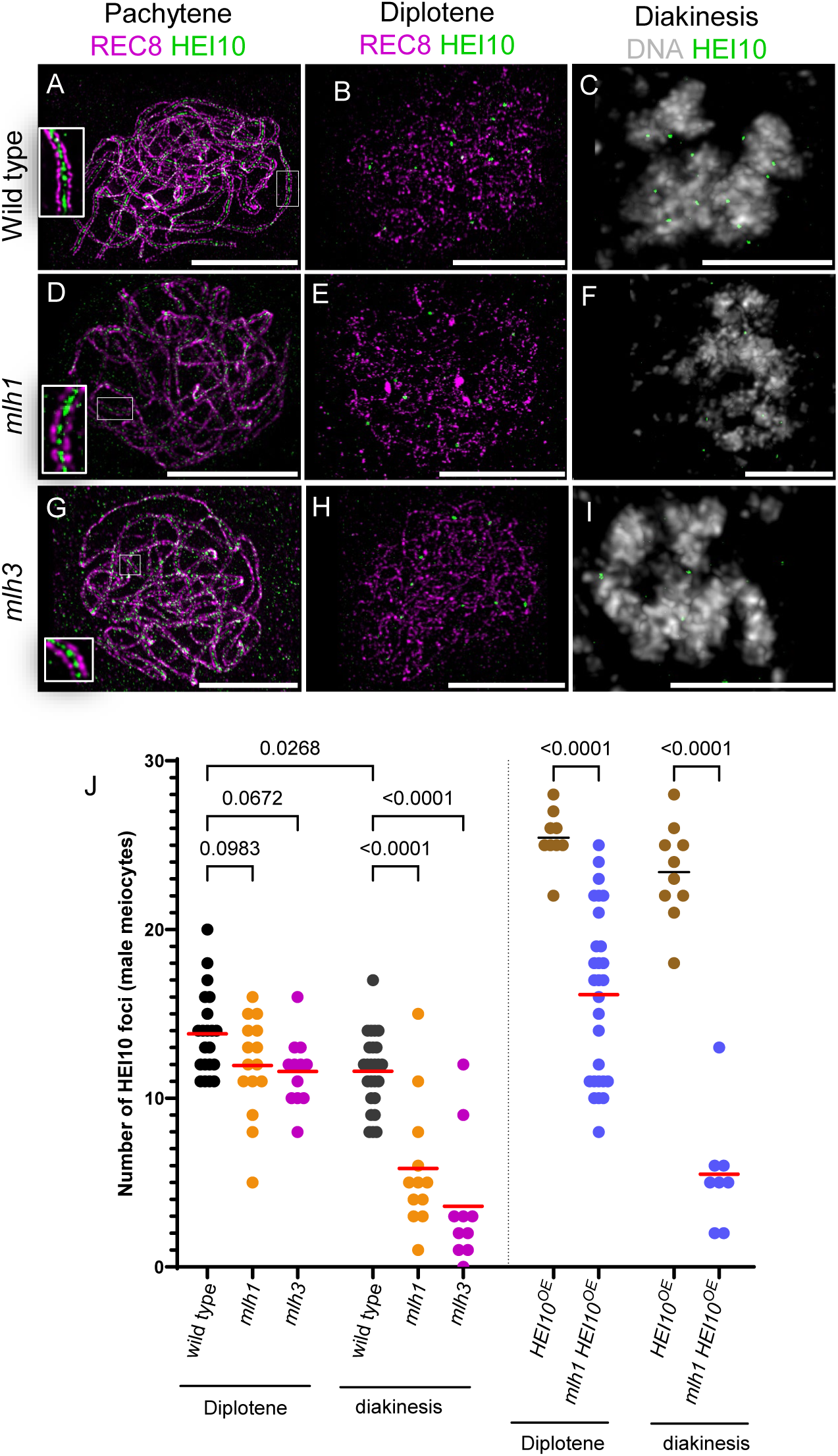
MLH1 and MLH3 Stabilize HEI10 foci. (A-I) Immunostaining of male meiocytes showing REC8 (purple) and HEI10 (green) and DNA (DAPI, grey) from pachytene to diakinesis in wild-type, *mlh1* and *mlh3* mutants. (J) Quantification of HEI10 foci in male meiocytes across genotypes at Diplotene and Diakinesis stages. Each dot represents a single cell, and the red bar indicates the mean. P values are one-way ANOVA with uncorrected Fisher’s least significance difference (LSD).

Overexpression of HEI10 (HEI10^oe^) in wild-type plants results in an increase in the number of HEI10 foci at diplotene and diakinesis, and an increase in CO frequency. (14,25,27) (Figure 3J). HEI10 overexpression in *mlh1* also increased the number of HEI10 foci at diplotene, although significantly less than when HEI10 is overexpressed in the wild type, which could reflect the role of MLH1 in the formation and/or the stability of HEI10 foci. However, in *mlh1* HEI10^oe^ diakinesis, the number of foci dropped to a very low level, further supporting the role of MLH1 in stabilizing HEI10 foci (Figure 3J). We propose that HEI10 accumulates and form foci at designated CO sites in both wild-type and *mutl*γ mutants, but that in the absence of MutLγ, HEI10 disengages from the recombination intermediates at late prophase.

### Genetic crossovers are differently reduced in *mlh1* and *HEI10* +/-

To explore the effect of *mlh1* on genetic recombination, we generated a novel *mlh1* mutant allele in a second strain (L*er*; *mlh1-3*) using CRISPR (Figure S1) and produced Col/L*er mlh1* hybrids by crossing. The *mlh1* hybrid had a meiotic defect similar to the *mlh1* inbred Col with an average of 1.1 univalents at metaphase I in male meiosis (Figure 4A) and reduced fertility (Figure 4B-C). Hybrids *mlh1* were backcrossed as female or male to wild-type Col, and the resulting progenies were sequenced to analyze recombination in both sexes separately, as previously described (14,24,28). In the wild type, the number of COs per transmitted gamete was substantially higher in males than in females (Figure 4D, 5.0 COs vs 3.3 COs, two-tailed Mann-Whitney test p<10^-6^), consistent with previous reports (28,29). In *mlh1*, the number of transmitted COs was significantly reduced to 3.6 in males (p<10^-6^) and to 2.6 in female meiosis (p=0.0002) (Figure 4D). Heterochiasmy is thus maintained in *mlh1* with ∼40% more COs in male than female meiosis. Note that these measures likely underestimate the CO defects (or overestimate the residual COs), as achiasmatic chromosomes have less chance to be transmitted. We also detected trisomies in both female (15/159, 9.4%) and male (3/124, 2.4%) populations, consistent with chromosome missegregations due to the presence of univalents at meiosis I.

**Figure 4:**
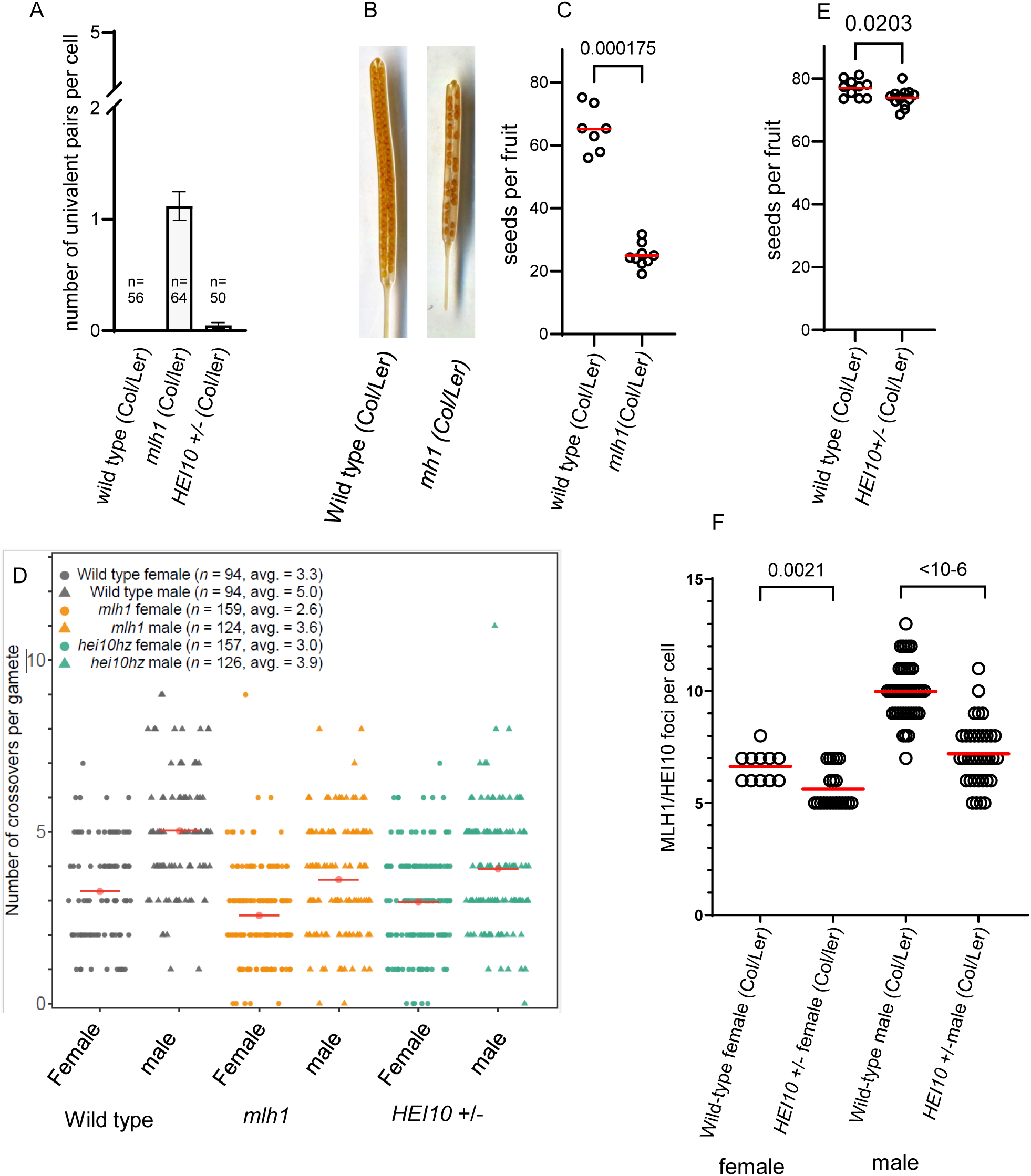
Genetic crossovers are differently affected in *mlh1* and *HEI10 +/-*, (A) Quantifying of the number of univalent pairs at metaphase I of male meiosis in Col/L*er* F1 hybrids of three different genotypes (wild-type *mlh1-/-* and *HEI10 +/-*). If no univalent pairs are observed, no bars are displayed. The number n of analyzed cells is indicated. Bars indicate mean +/-SEM. (B) Representative images of fruits from wild-type and *mlh1* mutant hybrid plants. (C) Quantification of fertility in *mlh1* hybrids. Each dot represents the average number of seeds per fruit (out of 10 fruits) for an individual plant. The red line indicates the mean number of seed per fruit for a given genotype. P values are Mann-Whitney tests. (D) Number of genetic crossovers per transmitted gamete in back-cross populations. Each dot corresponds to an individual female (circle) and male (triangle) gamete. Red lines indicate the mean CO number. The number of analyzed gametes and the average crossover genotype for each sex/genotype are shown. (E) Quantification of fertility in *HEI10 +/-* hybrids. Each dot represents the average number of seeds per fruit (out of 10 fruits) for an individual plant. The red line indicates the mean number of seed per fruit for a given genotype. Statistical significance was assessed using the Mann-Whitney test. (F) Quantification of HEI10/MLH1 co-foci in *HEI10 +/-* male meiocytes at diplotene and diakinesis stage. Each dot represents a cell, and the horizontal red bar indicates the mean. P values are Mann-Whitney test.

Based on the result described above, we propose that *mlh1* is defective in CO implementation but not in the designation of CO sites. We thought to compare recombination in *mlh1* to a context in which COs are reduced by a modification of the CO designation process. This is the case when HEI10 levels are reduced, leading to a reduction of COs in plants heterozygous for a defective *HEI10* allele (25,27). We produced *HEI10* +/- hybrids with a wild-type functional *HEI10* allele from L*er* and the *hei10-2* mutant allele from Col (15). In this *HEI10+/-* Col/L*er* hybrid context, the number of MLH1/HEI10 foci at diplotene was reduced compared to the corresponding wild type in both female and male meiocytes, consistent with a decrease in class I COs (Figure 4F). In contrast to *mlh1*, the number of univalents in *HEI10* +/- was very low, fertility was maintained at high levels (Figure 4A, 4E), and no aneuploidy was detected in progenies (0/126 and 0/157 for females and males, respectively). This suggests that the obligate CO is maintained in *HEI10* +/- despite a reduced number of COs. This contrasts with *mlh1* in which the obligate CO is lost while the number of COs designated site appears to be maintained (HEI10 foci at diplotene, Figure 3). Hybrids *HEI10 +/-* were then backcrossed as female and male to wild-type Col, and the resulting progenies were sequenced to analyze genetic recombination (Figure 4D). In females, the mean CO number is not significantly reduced (2.97 compared to 3.27 in wild type, p=0.15), likely reflecting that the number of COs is close to the minimum ensuring the obligate CO (5 COs per meiocyte = 2.5 COs per gamete). The number of COs in females is slightly lower in *mlh1* than in *hei10 +/-* (p=0.0049, Figure 4D). In *HEI10* +/- males, the number of COs was significantly reduced compared to the wild type (from 5 to 3.9, p=7.10^-6^), reaching similar levels than in *mlh1* males (p=0.18, Figure 4D). Beyond the reduction in CO numbers, the distribution of CO along chromosomes is not dramatically affected in *mlh1* and *hei10* +/- compared to wild type (Figure 5A, Figure S2).

**Figure 5:**
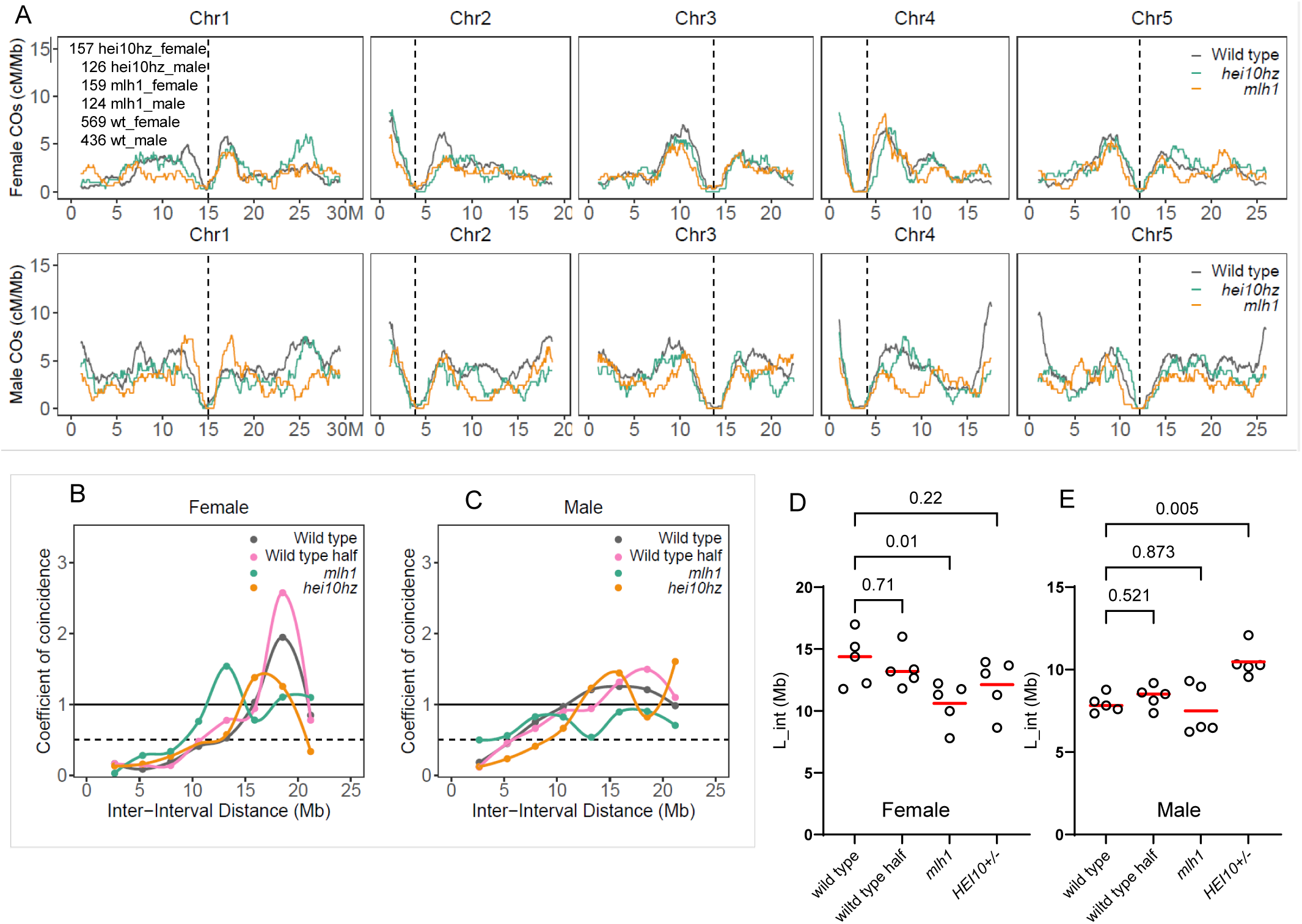
Crossover Distribution and Interference in *mlh1* and *HEI10* +/-. (A) Distribution of crossovers along chromosomes in females (top) and males (bottom) of wild-type, *mlh1* and *HEI10* +/-. The Y axis is CO frequency in centimorgan per Mb. The X-axis is the position in Mb (TAIR10). The vertical dashed line marks the position of the centromere. To gain power wild-type data from previous experiments were pooled with wild types from this study (Figure S2). (B) Coefficient of coincidence (CoC) curves for female and male meiosis. The CoC is displayed on the Y-axis, while the X-axis shows the inter-interval distance in Mb. A CoC value of 1 suggests independent CO occurrence, whereas a value near 0 indicates crossover interference. (D-E) Interference Length (L_int), which measures the shift in inter-CO distances due to interference. Higher L_int values indicate stronger crossover interference. Each dot is the L-int for an individual chromosome. Tests are Kruskal-Wallis and uncorrected Dunn’s test.

CO interference prevents the occurrence of close double-COs. It can be analyzed with the coefficient of coincidence (CoC), which divides the observed frequency of COs occurring concomitantly in two intervals by the expected frequency if COs were independent (product of the frequencies of COs in each interval). A CoC of 1 indicates that COs occur independently of one another while a CoC close to 0 reveals CO interference. A CoC curve is obtained by plotting the CoC values versus the distances separating the two considered intervals. In wild type, CoC curves are below 1 for short distances in both female and male meiosis, confirming the presence of CO interference (Figure 5B-C). When measured in the genomic space, interference is stronger in wild-type female meiosis, with the CoC curve reaching 1 at ∼16Mb, compared to males where the curve reaches 1 at shorter distances (∼11Mb). Another newly developed method to measure interference is L_int, which computes the shift in inter-CO distances due to interference (30). One advantage of L_int is that it provides a numerical value of interference, that we computed for each chromosome. In the wild type, L_int values are larger in female than male datasets, confirming the strongest CO interference in the genomic space (Figure 5D-E). This is consistent with previous observations and is attributed to shorter (µm) chromosome axis in females (with the same amount of DNA), leading to stronger interference in the genomic space (14,24,28,29). One intrinsic property of CO interference as measured by these two methods is the insensitivity to random sub-sampling of CO (30,31). Theoretically, losing randomly half of the COs is expected to maintain identical CoC curves and L_int values. We tested this property by randomly eliminating half of the COs in the wild-type dataset, and as expected the CoC curves and L_int were not significantly modified (Figure 5B-E).

In *hei10* +/- males, interference was increased in comparison to wild-type, with the CoC curve shifted to the right and increased L_int values (Figure 5B-E). This is consistent with previous cytological observations which concluded that decreased HEI10 levels increase interference and concomitantly reduce CO numbers, as predicted by the Coarsening Model (25). We conclude that in *hei10* +/- males, the decrease in COs is due to a modified process of CO designation. In *hei10* +/- females, the CoC curve and L_int were not significantly modified (Figure 5B, 5D), consistent with a nonsignificant decrease in CO number (Figure 4D), probably because interference is already very strong in wild-type females.

In *mlh1*, even if CO numbers are strongly reduced (more than in *hei10* +/-), the CO interference was not increased. The CoC curves and L_int values may suggest a slight decrease in CO interference, but this is at the edge of being significant. This is in clear contrast to *hei10* +/- males, where reduced CO number is associated with increased interference. This suggests that the designation process is not affected in *mlh1* and is consistent with the proposal that implementation is defective, with a fraction of the designated site randomly failing to produce COs, reducing CO number without affecting the interference values.

Altogether, we conclude that the function of MLH1/MLH3 is to impose designated recombination intermediates to be resolved exclusively as COs. In their absence, CO-designated intermediates (dHJs) are repaired by alternative enzymes, such as MUS81, that promote CO formation less efficiently, failing to ensure CO formation. These conclusions are compatible with the proposed role for MLH1/MLH3 in yeast and mammals -asymmetric resolution of dHJs exclusively as CO- and suggest that this function is widely conserved in eukaryotes. We cannot exclude an additional earlier role of MLH1 or MLH3 in the recombination process, but it would have no or a minor impact on the number of CO events (32,33). Finally, it should be noted that MLH1/MLH3 are not found in some species/lineages such as *Drosophila melanogaster* and *Caenorhabditis elegans*, suggesting they evolved alternative mechanisms to ensure CO maturation.

## Materials and methods

### Plant Materials and Growth Conditions

*Arabidopsis thaliana* plants were cultivated in growth chambers or greenhouses (21°C in 16 h day, 18 °C in 8h night, 60% humidity). Wild-type Col-0 and L*er*-1 are 186AV1B4 and 213AV1B1 from the Versailles *A.thaliana* stock center (http://publiclines.versailles.inra.fr/). The mutant alleles used in this study are: *mlh1-2* (Col, N1008089, SK25975, K#123), *mlh3-3* (Col, N619674, SALK_119674, K#309), *mus81-2* (Col; N607515, SALK_107515, K#133) (34), *msh5-3* (Col, N841758, SAIL_1056_F12, K#124), *HEI10^OE^* (Col, *C2 line,* pGreen0029-HEI10Col-H2, K#244) (27), *hei10-2* (Col, N514624, SALK_014624, K#010) (15), *mlh1-3* (L*er*, K#344) (Figure S1).

To generate double mutant *mlh1-2* HEI10^oe^ in Col background, homozygotes HEI10^oe^ (*C2*) were crossed as female with heterozygotes *mlh1-2* +/- as male. As the *MLH1* gene and the *HEI10* insertion are linked on chromosome 4, we grew the F3 population to isolate double homozygote plants. Those double *mlh1-2* HEI10^oe^ and wild-type control sister plants were used for MLH1/HEI10 foci counting. *mlh1-3* mutant allele in the L*er* background was obtained by CRISPR technology using two guides GATGATTACGGGAAAATCG and CCTGTGACTCCTCTGGTTG (35).

Transformations were performed with floral dipping. Plant transformants (T1) were selected by seed fluorescence. T2 seeds without fluorescence were selected and screened for mutations by PCR and Sanger sequencing of the targeted locus.

To generate *mlh1-2/mlh1-3* in Col/L*er* and the wild-type control Col/L*er*, heterozygotes *mlh1-2* +/- were crossed as female with heterozygotes *mlh1-3* +/- as male. The sister plants *mlh1* -/- and wild-type controls were used for HEI10 foci counting, bivalent analysis, and sterility analysis and were reciprocally backcrossed with wild-type Col to generate the populations to be sequenced.

To generate *HEI10* +/- in Col/L*er*, *hei10-2 +/-* from Col were crossed as female with L*er* as male. Sister plants *HEI10* +/- and wild-type were used for MLH1/HEI10 co-foci counting, bivalent analysis, and fertility measurements and were reciprocally backcrossed with wild-type Col to generate populations to be sequenced.

### Fertility

The fertility analysis of Arabidopsis plants was measured by counting the number of seeds per silique. At least 10 siliques sampled on the primary stem were analyzed per plant. Sister wild-type and mutant plants from segregating populations grown in the same environment were compared. The siliques were incubated in 70% ethanol. Once the siliques were translucent, they were imaged on a regular scanner. Seeds were counted using ZEN Software (Carl Zeiss Microscopy).

### Cytology

DAPI spreads (36). Young inflorescences were harvested from Arabidopsis plants and fixed in freshly prepared 3:1 ethanol:acetic acid. The fixative was replaced twice, and the fixed sampled stored at 4°C. Flower buds roughly 0.5mm in size were isolated from inflorescences and washed twice in water and once in citrate buffer (10mM tri-sodium-citrate, pH 4.5 with HCl), then incubated in a digestion mix (0.3% (w/v) Pectolyase Y-23 (MP Biomedicals), 0.3% (w/v) Driselase (Sigma), 0.3% (w/v) Cellulase Onozuka R10 (Duchefa), 0.1% sodium azide, in 10 mM citrate buffer) for 2 hours at 37°C, washed twice with water and kept on ice. Four to five digested buds were transferred to a clean slide and macerated with a bent dissection needle. Roughly 15uL of 60% acetic acid was added to the mixture and stirred gently at 45°C on a hotplate for 1 minute. Another drop of 15ul of 60% glacial acetic acid was added again to the mixture and stirred for another 1 minute. The slide was then flushed with ice-cold 3:1 fixative first around the droplet and then directly. Slides were left to dry tilted at room temperature, then 10uL of mounting media with 2μg/mL DAPI was applied to the slide and a coverslip was added. Chromosomes were imaged with a Zeiss Axio Observer epifluorescence microscope.

Immunolocalizations were performed on cells with preserved 3-dimensional structures (37). For male meiocytes, Sepals and petals were removed from 0.35–0.45mm flower buds and collected in buffer A (KCl 80mM, NaCl 20mM, Pipes-NaOH 15mM, EGTA 0.5mM, EDTA 2mM, Sorbitol 80mM, DTT 1mM, Spermine 0.15mM, spermidine 0.5mM). For female meiocytes, 0.8–1.2mm pistils were collected and their stigmata were cut off and collected in buffer A. Male and female samples were fixed by incubation in bufferA+2% formaldehyde for 30 mins under vacuum. Buds or pistils were then washed in buffer A for 10 minutes and digested at 37° for 1 hour (0.3% cellulase, 0.3% pectolyase Y23, 0.3% driselase, 0.1% sodium azide in citrate buffer). After a wash in buffer A, digested buds or pistils were kept in buffer A on ice. To make the embedding, 5-8 buds or pistils were placed in 6µl of buffer A on a 18mmx18mm High Precision coverslip, and anthers or pistils were dissected and squashed to extrude meiocytes. A 3µl drop of activated polyacrylamide solution (25µl 15% polyacrylamide (SIGMA A3574) in buffer A + 1.25µl of 20% sodium sulfite + 1.25µl of 20% ammonium persulfate) is added to the meiocytes and a second coverslip is placed on the top, with gentle pressure. The polyacrylamide gels were left to polymerize for 1h and then the two coverslips were separated. The coverslips covered by a gel pad were incubated in 1X PBS, 1% Triton X-100, 1mM EDTA for 1h with agitation, followed by 2h in blocking buffer (3% BSA in 1X PBS + 0.1% Tween 20) at room temperature. Coverslips were then incubated with 100µl of primary antibody in blocking buffer at 4°C in a humid chamber for 48h. Coverslips were washed four times 30min with 1X PBS, 0.1% Triton X-100. One hundred microliters of the appropriate fluorophore-conjugated secondary antibodies in blocking buffer were applied (1:250) and incubated at room temperature overnight in the dark. Gels were washed four times 20 min with 1X PBS, 0.1% Triton X-100. 6µl of SlowFade™ Gold (for super resolution microscopy) + 10µM DAPI were used for mounting the coverslip with a slide, that was sealed with nail polish. The primary antibodies used were: anti-REC8 raised in rat (38) (lab code PAK036, dilution 1:250), anti-MLH1 raised in rabbit (12) (PAK017, 1:200), anti-HEI10 raised in chicken (15) (PAK046, 1:5000). Secondary antibodies were Abberior StarRed, StarOrange and STARgreen for STED microscopy. Images for MLH1/HEI10 co-foci analysis were taken with the Abberior instrument facility line (www.abberior-instruments.com) using 561nm and 640nm excitation lasers (for STAR Orange and STAR Red, respectively) and a 775nm STED depletion laser. Confocal images were taken with the same instrument with a 485nm excitation laser (for Stargreen). Images were deconvolved with Huygens Essential version 20.04 (Scientific Volume Imaging, The Netherlands, http://svi.nl), using the CMLE algorithm, with lateral drift stabilization; SNR: 7 for STED images, 20 for confocal images; 40 iterations; and quality threshold: 0.5. Maximum intensity projections and contrast adjustments were done with Huygens Essential.

### Genetic crossover and aneuploidy analysis by sequencing

Populations to be sequenced were grown in the greenhouse for three weeks (16-h day/8-h night) and four days in the dark. Leaf samples (100–150 mg) were used for DNA extraction and library preparation for Illumina sequencing (39) at the Max Planck Genome Center, Cologne, Germany (https://mpgc.mpipz.mpg.de/home/). To create a list of high-quality SNP markers, a strategy of combining the whole-genome alignment short-read mapping was used. The raw sequencing data was aligned to the TAIR10 Col reference genome (40) by BWA v0.7.17-r1188 (41) with the default parameters. COs were detected using a sliding window-based method, with a window size of 50 kb and a sliding step of 25 kb (14,24,28). The identified COs were also manually and randomly checked by using inGAP-family (42).

A total of 157 and 226 COs were detected from the 47 female and 47 male wild-type plants (sister plants of *mlh1*), 408 and 448 COs were detected from the 159 female and 124 male *mlh1* plants, 150 and 247 COs were detected from the 47 female and 47 male wild-type plants (sister plants of *hei10* +/-), 466 and 495 COs were detected from the 157 female and 126 male *hei10* +/- plants, respectively. The list of 1,192 and 1,587 COs of the wild-type female and male populations (428 and 294 plants, ArrayExpress number E-MTAB-11254) (28) and the list of 138 and 254 COs of the wild-type female and male populations (47 and 48 plants, ArrayExpress number E-MTAB-12838) (43) from the previous studies were used for the analysis of CO distribution and CO interference.

To analyze the CO interference, we calculated the Coefficient of Coincidence (CoC) with (44) and L_int values (30). The sequencing depths of each chromosome were evaluated by Mosdepth v0.2.7 (14,45), with a window size of 100 kb. The sample with more than 1.2-fold difference in sequencing depths between chromosomes was considered aneuploid.

## Data availability

The number and list of identified COs in the female and male populations of wild type, *mlh1*, and *hei10* +/- can be accessed in Supplementary Table 1-2. The raw sequencing data generated in this study have been deposited in the ArrayExpress EMBL-EBI database under accession codes E-MTAB-14422 and E-MTAB-14423.

## Acknowledgments

This work was supported by core funding from the Max Planck Society, and Alexander von Humboldt Fellowships to Q.L. and J.B.F. We thank Ian Henderson for kindly providing the HEI10 C2 line.

