## Supplemental figures for "MutLγ enforces meiotic crossovers in *Arabidopsis thaliana*"

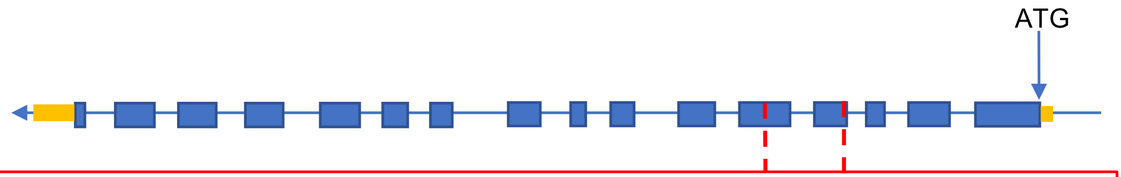


*MLH1*-exon4-5
GTGGAGAATTTGTTCTACAATATGATTGCTAGAAGGAAGACACTTCAAAATTCTGCTGATGATTACGGGAAAAT
CGTGGATTTGCTGAGCCGGATGGCTATTCATTACAATAATGTCAGCTTTTCTTGTCGAAAGgttggtatacttattaactaa
actagtcgtatttagcaatcttttgtgttttggaattgtgaagtgaattttggatgatactgtctcgccagCATGGAGCTGTTAAGGCTGATGTTCACT
CAGTCGTGTCACCTTCAAGGCTTGATTCAATTAGGTCTGTATATGGAGTATCAGTTGCAAAGAACTTGATGAAA
GTAGAAGTTTCCTCCTGTGACTCCTCTGGTTGTACTTTTGATATGGAGGGTTTCATATCCAATTCTAACTATGTT
GCTAAGAAGACTATATTGGTGCTTTTCATTAATG

Protein sequence:
MIDDSSLTAEMEEEESPATTIVPREPPKIQRLEESVVNRIAAGEVIQRPVSAVKELVENSLDAD
SSSISVVVKDGGLKLIQVSDDGHGIRREDLPILCERHTTSKLTKFEDLFSLSSMGFRGEALASM
TYVAHVTVTTITKGQIHGYRVSYRDGVMEHEPKACAAVKGTQIMVENLFYNMIARRKTLQNS
ADDYGKIVDLLSRMAIHYNNVSFSCRKHGAVKADVHSVVSPSRLDSIRSVYGVSVAKNLMKV
EVSSCDSSGCTFDMEGFISNSNYVAKKTILVLFINDRLVECSALKRAIEIVYAATLPKASKPFVY
MSINLPREHVDINIHPTKKEVSLLNQEIIIEMIQSEVEVKLRNANDTRTFQEQKVEYIQSTLTSQK
SDSPVSQKPSGQKTQKVPVNKMVRTDSSDPAGRLHAFLQPKPQSLPDKVSSLSVVRSSVRQ
RRNPKETADLSSVQELIAGVDSCCHPGMLETVRNCTYVGMADDVFALVQYNTHLYLANVVN
LSKELMYQQTLRRFAHFNAIQLSDPAPLSELILLALKEEDLDPGNDTKDDLKERIAEMNTELLK
EKAEMLEEYFSVHIDSSANLSRLPVILDQYTPDMDRVPEFLLCLGNDVEWEDEKSCFQGVSA
AIGNFYAMHPPLLPNPSGDGIQFYSKRGESSQEKSDLEGNVDMEDNLDQDLLSDAENAWAQ
REWSIQHVLFPSMRLFLKPPASMASNGTFVKVASLEKLYKIFERC*

**Figure S1: MLH1 gene and *mlh1-3* mutation.**

The structure of the MLH1 gene is shown, with the coding sequence in blue and UTRs in yellow. The 326pb deletion in the genomic sequence of L*er* *mlh1-3* mutant is represented in red. The underlined sequence corresponds to the position of the used CRISPR guide. The in-frame deletion predicts a deletion of 78 amino acids in the MLH1 protein sequence (M179 to C256), deleting a large part of the MUTL domain (interpro038973 29-401).

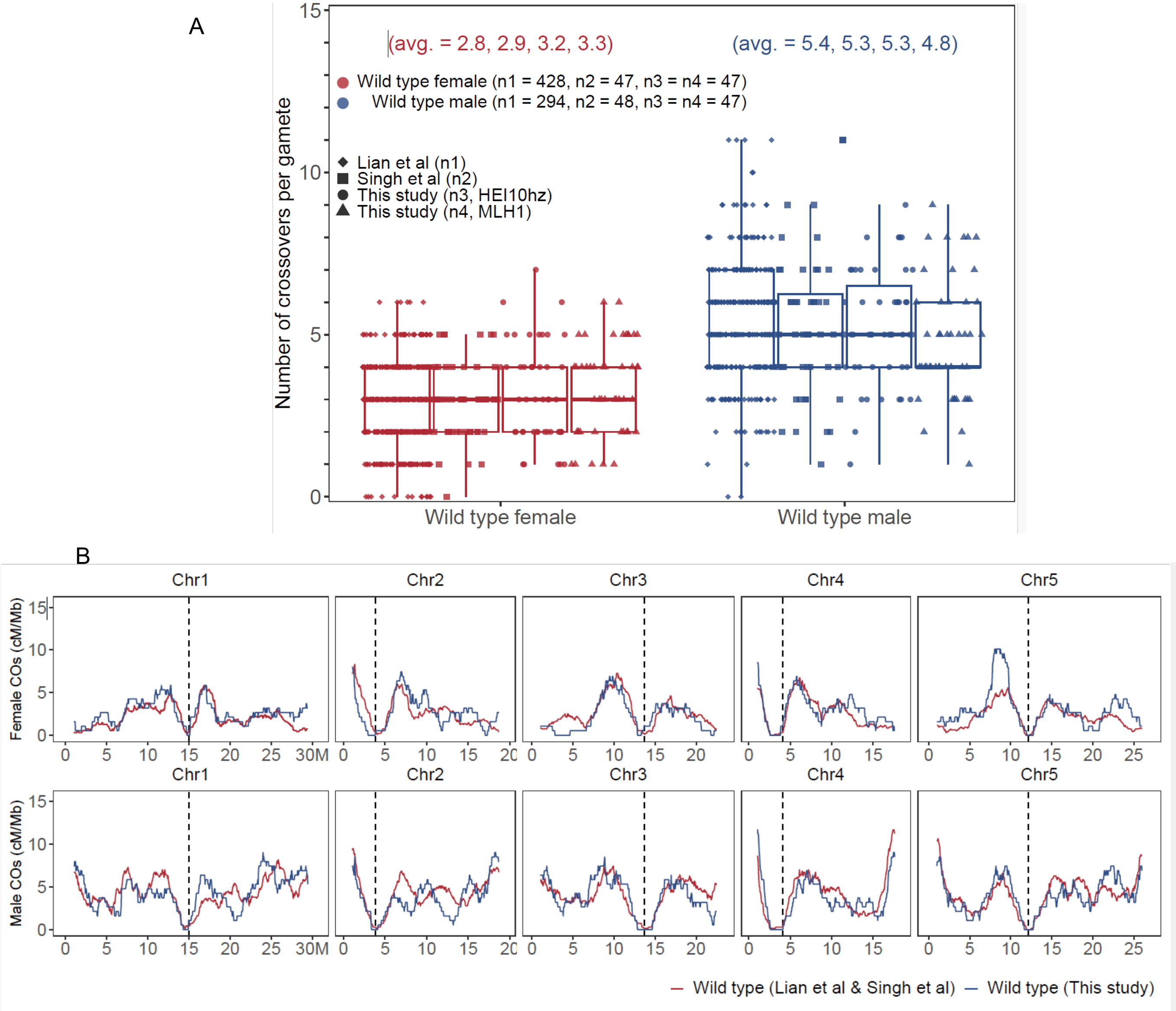


**Figure S2: Crossover number and distribution in wild-type datasets.**
(A) Number of crossovers per transmitted female (red) and male (blue) gametes, in Col/L*er* wild-types from independent experiments. Each dot corresponds to an individual plant. The four populations are from Lian et al. (Lian, Solier et al. 2022), wild-type controls from Singh et al. (Singh, Lian et al. 2023), control sister plants from *HEI10 +/-* (This study) and control sister plants from *mlh1* (This study). (B) Distribution of crossovers (COs) in females and males. The blue line shows the wild type from this study (merge controls of *HEI10* +/- and *mlh1.* n=94 females and 94 males). The red line shows wild-type from previous work (merge wild-type from Lian et al., and wild-type control of *heip1* from Singh et al. Total n=475 for females, and n=342 for males).
